# Ambient temperature and genotype differentially affect developmental and phenotypic plasticity in *Arabidopsis thaliana*

**DOI:** 10.1101/017285

**Authors:** Carla Ibañez, Yvonne Poeschl, Tom Peterson, Julia Bellstädt, Kathrin Denk, Andreas Gogol-Döring, Marcel Quint, Carolin Delker

**Author notes:** Co-first authors (Carla Ibañez, Yvonne Poeschl).

## Abstract

**Background:** Global increase in ambient temperatures constitute a significant challenge to wild and cultivated plant species. Forward genetic analyses of individual temperature-responsive traits have resulted in the identification of several signaling and response components. However, a comprehensive knowledge about temperature sensitivity of different developmental stages and the contribution of natural variation is still scarce and fragmented at best.

**Results:** Here, we systematically analyze thermomorphogenesis throughout a complete life cycle in ten natural *Arabidopsis thaliana* accessions grown in four different temperatures ranging from 16 to 28 °C. We used Q_10_, GxE, phenotypic divergence and correlation analyses to assess temperature sensitivity and genotype effects of more than 30 morphometric and developmental traits representing five phenotype classes. We found that genotype and temperature differentially affected plant growth and development with variing strengths. Furthermore, overall correlations among phenotypic temperature responses was relatively low which seems to be caused by differential capacities for temperature adaptations of individual accessions.

**Conclusion:** Genotype-specific temperature responses may be attractive targets for future forward genetic approaches and accession-specific thermomorphogenesis maps may aid the assessment of functional relevance of known and novel regulatory components.

## Background

Recurrent changes in ambient temperature provide plants with essential information about time of day and seasons. Yet, even small changes in mean ambient temperatures can profoundly affect plant growth and development resulting in thermomorphogenic changes of plant architecture [1]. In crops like rice, a season-specific increase in the mean minimum temperature of 1 °C results in a ~10 % reduction in grain yield [2]. Likewise, up to 10 % of the yield stagnation of wheat and barley in Europe over the past two decades can be attributed to climate change [3]. Current projections indicate that mean global air temperatures will increase up to 4.8 °C by the end of the century [4,5]. Global warming will thus have significant implications on biodiversity and future food security.

Elevated ambient temperatures affect of course also wild species in their natural habitats. Long-term phenology studies of diverse plant populations have revealed an advance in first and peak flowering and alterations in the total length of flowering times [6,7]. Furthermore, estimates project that temperature effects alone will account for the extinction of up to one-third of all European plant species [8]. As the impact of changes in ambient temperature on crop plants and natural habitats emerge, a comprehensive understanding of themperature-mediated growth responses throughout development becomes paramount.

Our present knowledge on molecular responses to ambient temperature changes has significantly progressed by studies in *Arabidopsis thaliana.* Model thermomorphogenesis phenotypes such as hypocotyl elongation [9], hyponastic leaf movement [10], and alterations in flowering time have served in various genetic approaches to identify relevant molecular players (reviewed in [1]. In this regard, exploiting naturally occurring genetic variation in these model traits has served as a valuable tool [11–16]. Primary signaling genes/proteins seem to function in response to both temperature and light stimuli. Prominent members of this network are photoreceptors such as CRYPTOCHROME 1 (CRY1 [17]), or the recently identified thermosensor PHYTOCHROME B [18,19]. Further components include PHYTOCHROME INTERACTING FACTOR 4 (PIF4, [20–22], DE-ETIOLATED 1, CONSTITUTIVELY PHOTOMORPHOGENIC 1, ELONGATED-HYPOCOTYL 5 [23–25] and EARLY FLOWERING 3 (ELF3); the latter as a component of the circadian clock [12,13].

The investigation of signaling pathways that translate temperature stimuli into qualitative and quantitative developmental responses has so far largely been limited to either seedling development or flowering time. However, it seems likely that temperature responses in different phases of development either require variations of a canonical signaling pathway or involve at least partially specific signaling components. To enable the dissection of thermomorphogenic signaling at different developmental stages, it is vital to gather a comprehensive understanding of the diversity of temperature reactions throughout plant development.

According to basic principles of thermodynamics, temperature-induced changes in free energy will affect the rates of biochemical reactions. As these effects should occur generally, albeit to different magnitudes, non-selective phenotypic responses can be expected to occur robustly and rather independently of genetic variation. Such traits may therefore be indicative of passive, thermodynamic effects on a multitude of processes. Alternatively, robust temperature responses may be due to thermodynamic effects on highly conserved signaling elements. These may be attractive targets for classic mutagenesis screens to identify the relevant regulatory components. In contrast, natural variation in thermomorphogenesis traits is likely the consequence of variability in one or several specific signaling or response components. It may be addressed by quantitative genetic approaches to identify regulators that contribute to variable temperature responses. Such genes may represent attractive candidates for targeted breeding approaches.

In this study we aim to (i) provide a map of developmental phenotypes that are sensitive to ambient temperature effects throughout a life cycle in the model organism *A. thaliana,* (ii) identify traits that are robustly affected by temperature with little variation among different accessions, and ask (iii) which traits are affected differentially by different genotypes and thus show natural variation in temperature responses.

To realize this, we performed a profiling of numerous developmental and morphological traits which can be sorted into five main categories: juvenile vegetative stage, adult vegetative stage, reproductive stage, morphometric parameters and yield-associated traits. Phenotypes were analyzed in a subset of ten *A. thaliana* accessions which were grown at 16, 20, 24, and 28 °C in climate-controlled environments. Knowing that even a small randomly selected set of *A. thaliana* accessions covers a wide spectrum of genetic diversity [26], we chose to analyze commonly used lab accessions such as Col-0, Ler-1 and Ws-2, accessions known to react hypersensitively to elevated temperature (e.g., Rrs-7, [24,27], and parental lines of available mapping populations such as Bay-0, Sha, and Cvi-0.

In addition to a meta-analysis of the phenotypic data, we provide accession-specific developmental reference maps of temperature responses that can serve as resources for future experimental approaches in the analysis of ambient temperature responses in *A. thaliana.*

## Materials and Methods

### Plant material and growth conditions

Phenotypic parameters (Fig. 1) were assessed in *A. thaliana* accessions that were obtained from the Nottingham Arabidopsis Stock Centre [28]. Morphological markers and time points of analyses are described in Additional file 1. Detailed information on stock numbers and geographic origin of Arabidopsis accessions are listed in Additional file 2. For seedling stage analyses, surface-sterilized seeds were stratified for 3 days in deionized water at 4 °C and subsequently placed on *A. thaliana* solution (ATS) nutrient medium [29]. Seeds were germinated and cultivated in climate-controlled growth cabinets (Percival, AR66-L2) at constant temperatures of 16, 20, 24 or 28 °C under long day photoperiods (16h light/8h dark) and a photosynthetically active fluence rate (PAR) of 90 μmol·m^−2^·sec^−1^ of cool white fluorescent lamps. We refrained from including a vernalization step because the primary focus of this study was to record morphology and development in response to different constant ambient temperature conditions.

**Figure 1:**
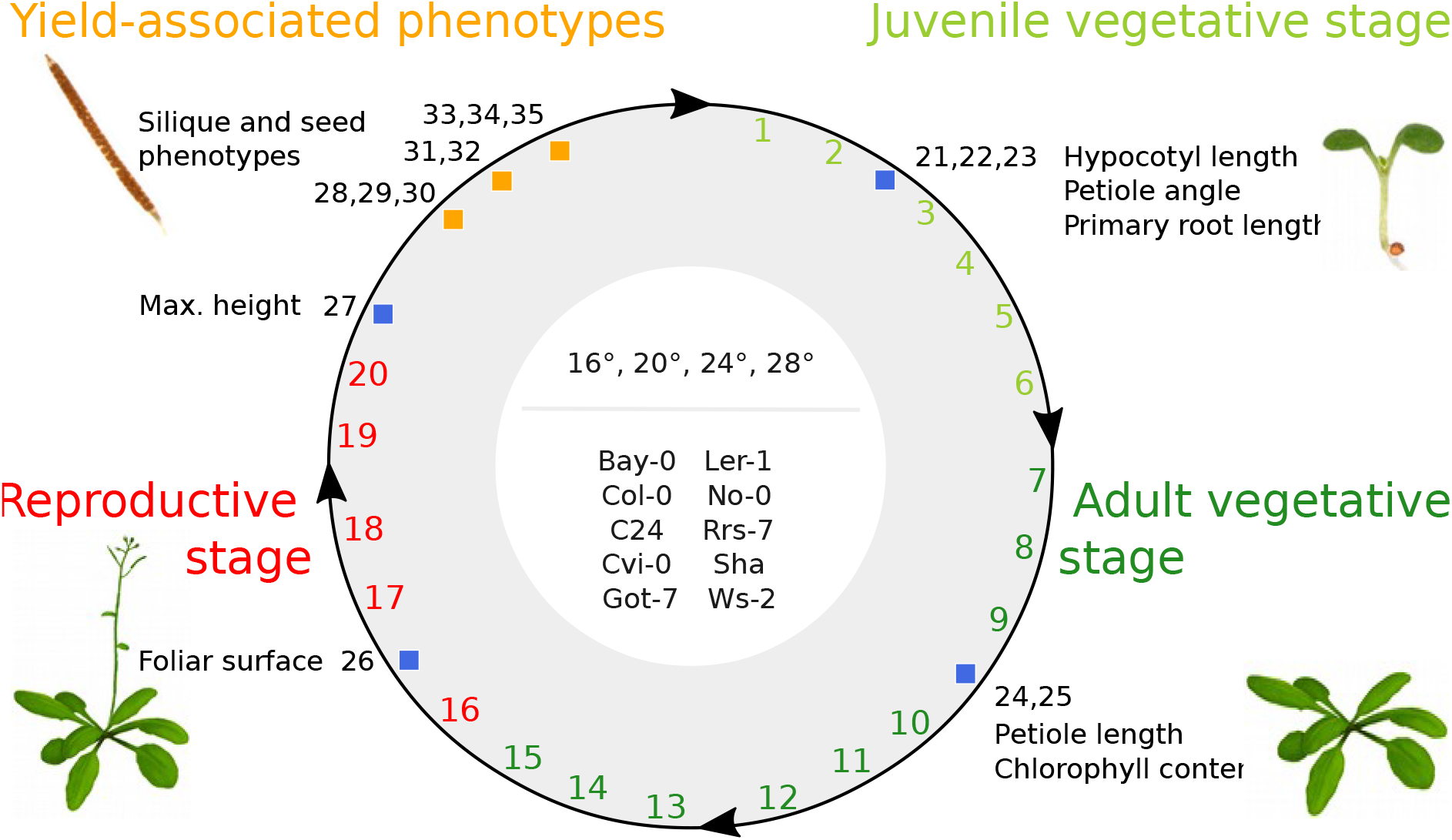
Phenotypic profiling approach. Schematic representation of the accessions, cultivation temperatures (°C) and phenotype classes used in the phenotypic profiling approach. Numbers indicate individual traits listed in Additional file 16 and are color-coded according to the corresponding phenotype class. Blue and orange squares indicate phenotypes sorted into *'morphometric phenotypes'* and *'yield-associated phenotype'* classes, respectively. Their position is indicative for the developmental stage at time of assessment.

Germination rates were assessed daily and hypocotyl, root length, and petiole angles were measured in 7 days old seedlings (n > 15) with ImageJ [30] and Root Detection [31].

All other analyses were performed on soil-grown plants cultivated in growth cabinets (Percival) at a PAR of 140 μmol·m^−2^·sec^−1^ and long day photoperiods (16h light/8h dark). After imbibition for 3 days at 4 °C, seeds were grown in individual 5 x 5 cm pots, which were randomized twice a week to minimize position effects. Relative humidity of growth cabinets was maintained at 70 % and plants were watered by subirrigation. Plants (n > 15) were photographed daily for subsequent determination of phenotypic parameters (leaf number, rosette area and petiole length) using Image J (http://imagej.nih.gov/ij/). Determination of developmental progression largely followed the stages defined in Boyes et al. [32]. The vegetative growth period was divided in a juvenile phase (germination to initiation of the fifth rosette leave) and an adult vegetative stage (initiation of the sixth rosette leave to floral transition). At transition to the reproductive growth phase, the number of leaves was determined by manual counting in addition to recording the number of days after germination. Spectrophotometric determination of chlorophyll content was performed as described in [33].

### Data analysis

Visualization and statistical analyses of the data were performed using the software R [34]. Box plots were generated using the *boxplot* function contained in the graphics i package. Heat maps were generated using the *heatmap.2* function contained in the gplots package.

ANOVAs for a single factor (either accession or temperature) and Tukey’s ‘Honest Significant Difference’ test as *post hoc* test were performed using the *anova* and *TukeyHSD* function, respectively, which are both contained in the R stats package.

Variation in phenotype expression was analyzed by 2-way ANOVA according to Nicotra [35] and Whitman and Agrawal [36] to test each phenotype for a significant effect of genotype (G, accession) or environment (*E*, temperature), and a significant genotype by environment interaction (GxE). Reaction norms for each analysis are shown in Additional file 3.

### Q_10_ temperature coefficient

The Q_10_ temperature coefficient was calculated according to Loveys [37]

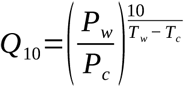

where P_w_ and P_c_ are the trait values at the warmer and cooler temperatures, respectively. T_w_ and T_c_ represent the corresponding temperatures in °C. We computed the geometric mean of the six Q_10_ values of all pairwise temperature combinations for each phenotypic trait to avoid artifacts caused by differential reaction norms/response shapes.

### Index of phenotypic divergence (P_st_)

Calculation of the index of phenotypic divergence (P_st_ [38,39]) as a measure to quantify variation in each phenotypic trait was calculated as previously described by Storz [38] as

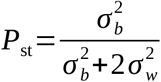

where *σ_b_*^2^ is the variance between populations, and *σ_w_*^2^ is the variance within populations. The ANOVA framework was used to partition the variances to get unbiased estimates for *σ_b_*^2^ and *σ_w_*^2^.

Using the two factorial design, two types of indices of phenotypic variation of a trait/phenotype were considered separately. The index of phenotypic divergence for genotypes (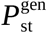) at a defined temperature level can be computed to measure the effect/impact of the genotype on the variation whereas the index of phenotypic divergence for temperatures (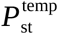) provides a measure for the effect of temperature on the observed variation for individual genotypes.

### Principal component analysis (PCA)

Arithmetic means for each genotype-temperature pair were computed except for six traits (germination, 13 rosette leaves, 14 rosette leaves, silique production, chlorophyl content (a+b), and foliar surface) due to too many missing values. The remaining 28 traits contained at most eight missing values (randomly distributed).which were replaced per trait by the arithmetic mean of the respective trait values. PCA was perfomed using the *prcomp* function contained in the R stats package. Due to the different units and scales of the traits the data was not only to centered but also to scaled by *prcomp.*

### Pairwise correlation analysis of traits

Trait values for rosette leave traits were summarized by arithmetic means to trait groups labeled *Juvenile vegetative stage (2–5 rosette leaves)* and *Adult vegetative stage (6–14 rosette leaves),* respectively. Similarly, *Inflorescence emergence, Flowering time_days* and *Flowering time_first flower* were combined to form the trait group *Flowering time (days)*. Spearman correlation coefficients were computed using the R stats package. Additionally, p values for each Spearman correlation coefficient were computed using the *cor.test* function. P values were subsequently corrected for multiple testing using the Benjamini-Hochberg correction implemented in the multtest package.

## Results

To assess phenotypic plasticity in a range of ambient temperatures, *A. thaliana* plants were cultivated in parallel throughout an entire life cycle at four different temperatures (16, 20, 24 and 28 °C) under otherwise similar growth conditions (see Materials and methods for further details). More than 30 morphological and developmental traits were recorded representing the following five phenotype classes: juvenile vegetative, adult vegetative, and reproductive stages as well as morphometric and yield-associated phenotypes (Fig. 1 and Additional file 1).

### Temperature responses in the A. thaliana reference accession Col-0

In Col-0, almost all phenotypes analyzed in this study were affected by the cultivation in different ambient temperatures. Only seed weight and maximum height remained constant regardless of the growth temperature (Fig. 2a, Additional file 4). Among the temperature-sensitive traits were several growth-associated phenotypes in the juvenile vegetative stage. Primary root length, hypocotyl and petiole elongation all increased with elevated temperatures which concurs with previously published data [9,10]. As another example, yield-related traits, such as the number of siliques per plant and the number of seeds per silique decreased with an increase in ambient temperature (Fig. 2a).

**Figure 2:**
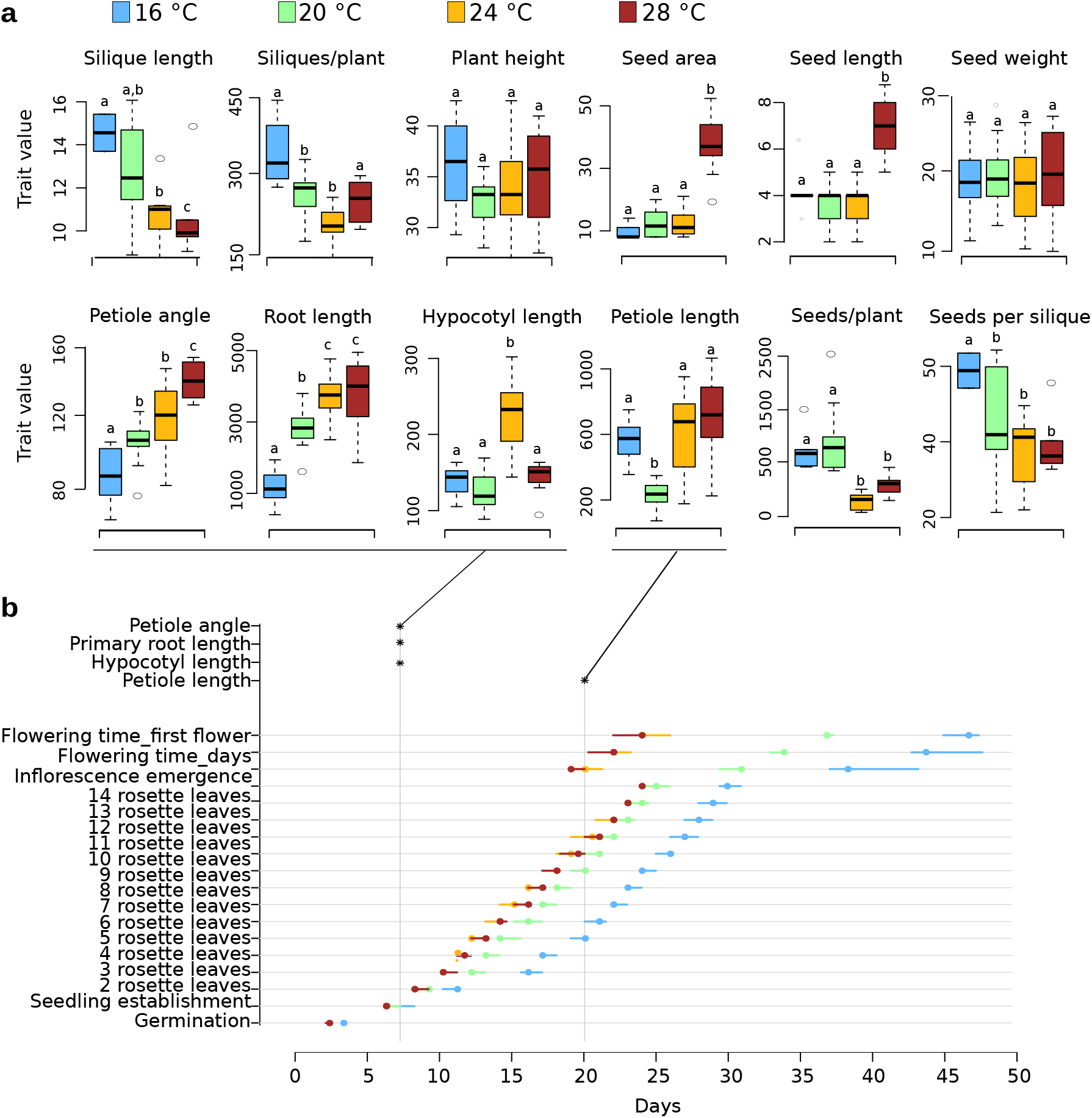
Col-0 growth and development in response to different ambient temperatures. (a) Quantification of phenotypic traits recorded at different growth temperatures. Box plots show median and interquartile ranges (IQR), outliers (> 1.5 times IQR) are shown as circles. Units for each trait are specified in Additional file 16. Different letters denote statistical differences (P > 0.05) among samples as assessed by one-factorial ANOVA and Tukey HSD. (b) Summary of temperature effects on developmental timing. Circles denote medians, bars denote IQRs (n > 15). Times of phenotypic assessment for selected traits in (a) are indicated by asterisks.

As reported previously, Col-0 plants showed a decrease in developmental time until flowering with increasing ambient temperatures [11]. The transition from the vegetative to the reproductive phase at 28 °C occurred about 25 days earlier than at 16 °C (Fig. 2a). Similarly, the number of rosette leaves developed at time of bolting differed by approximately 26 leaves between 28 °C and 16 °C (Additional file 4b).

The observation that only a very limited number of phenotypes were insensitive to cultivation in different temperatures clearly illustrates the fundamental impact of ambient temperature on plant growth and development.

### Natural variation of temperature responses

To assess whether the observed temperature responses in Col-0 are robust among *A. thaliana* accessions or which of the responses may be affected by natural variation, phenotypic profiling was performed in nine additional *A. thaliana* accessions parallel to the analysis in Col-0 (Additional files 4–13). Naturally, a panel of ten accessions does not comprehensively represent the world-wide gene pool of *A. thaliana.* However, it can be expected that even 10 randomly chosen natural accessions represent ~70 % of the allelic diversity in the *A. thaliana* gene pool [26]. Hence, the general assessment of thermo-responsive development in *A. thaliana* as well as the identification and discrimination between traits that generally seem to exhibit natural variation and those that may be genetically fixed within the gene pool is a realistic aim even with a set of 10 selected accessions.

To approximate and to compare temperature sensitivity of traits among different accessions, we calculated Q_10_ values for each individual trait and phenotype class for each analyzed genotype [37]. The Q_10_ quotient represents the factor by which a trait value changes if the ambient temperature increases by 10 °C. We calculated geometric means of all possible pairwise combinations of temperatures to minimize effects potentially caused by different response curves and used the log_2_Q_10_ for visualization as to retain high resolution in the presentation of the data.

Similarly to the response observed in Col-0 (Fig. 2), all analyzed genotypes showed a temperature-induced acceleration of vegetative development as indicated by negative log_2_Q_10_ values with low variability among accessions (Fig. 3a, b, Additional files 4–13). Considerably higher variation was observed in log_2_Q_10_ values of traits related to reproductive stages. As all accessions investigated were principally able to flower despite the lack of an extended cold period, none of them strictly required a vernalization treatment to transition to the reproductive phase. In contrast to the other accessions, Got-7 and Rrs-7, however, showed a significant delay in flowering time with increasing temperature (Fig. 3b). Got-7, for example, did not flower within the first 90 days of cultivation when grown in 24 or 28 °C. Thus, initiated leaf senescence at bolting stage prevented accurate determination of leaf number at the onset of flowering.

**Figure 3:**
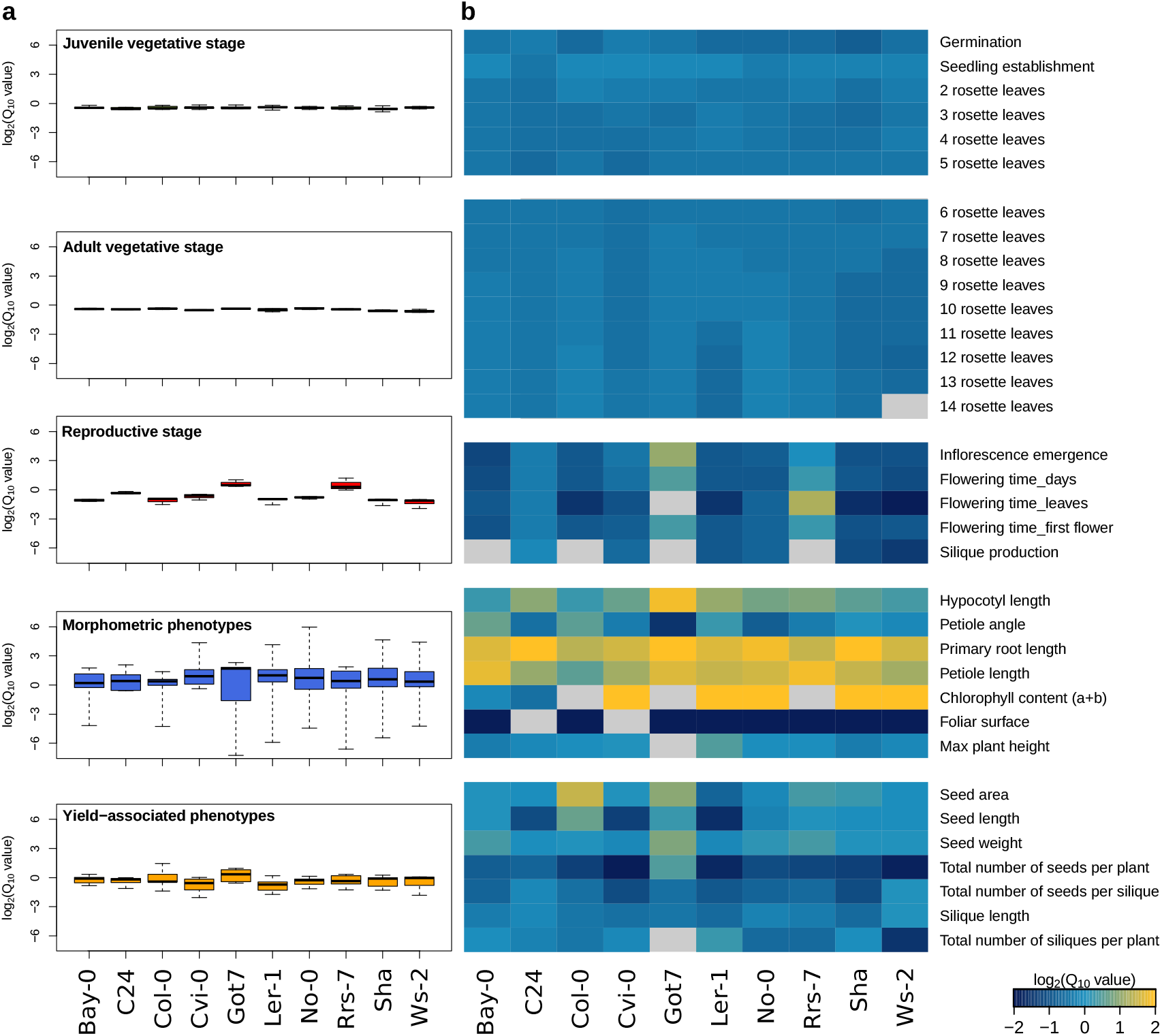
Natural variation in temperature sensitivity of phenotypic traits (Q_10_) Mean log_2_Q_10_ values for each accession (a) summarized in box plots for each phenotype class and (b) presented as a heatmap for all individual phenotypes. (a) Box plots show median and interquartile ranges (IQR), whiskers range from min. to max. values. (b) positive (increasing) and negative (decreasing) log_2_Q_10_ values are shown in yellow and blue, respectively with a log_2_Q_10_ cut-off value of 2 for better resolution. Missing data are denoted in light gray.

A direct comparison of leaf number and time of development further corroborates a sudden increase in variation at the transition to flowering (Additional file 14). However, at 16 °C and 20 °C several accessions contribute to the overall variability in the graph, whereas at 24 °C and 28 °C, C24 and Rrs-7 are the main determinants of variation due to their massive number of leaves corresponding to an extension of the vegetative growth phase (Additional file 14). Got-7 likely would increase this variation at 24 and 28 °C, but is missing in this representation due to the lack of flowering transition within 90 days. Here, the lack of vernalization may at least partially be a significant factor because cold treatment is explicitly recommended to induce earlier flowering for several Got-7 lines available at NASC/ABRC [40]. Natural variation in regulators such as FLM may contribute to this phenotype. However, as all accessions were able to flower at temperatures of 16 and 20 °C vernalization does not seem to be an essential requirement.

Taken together, juvenile and adult vegetative development remained highly conserved, whereas the reproductive stage and yield-associated traits showed higher variation between accessions and within individual accession, as indicated by the ranges/dimensions of the box plots in Fig. 3a. Here, high variation within a phenotype class indicates that temperature effects on individual traits within that class are highly variable. The strongest variation within accessions was observed for morphometric phenotypes such as hypocotyl and petiole length. In contrast, a high variation between accessions is indicative for differential responses of different genotypes which was most prominent in reproductive stage traits.

The differential variances of log_2_Q_10_ values among the two vegetative and the other phenotype classes indicated that genotype and environment effects may contribute differentially to phenotypic plasticity of different traits. We first used a 2-factorial ANOVA to assess which phenotypes show significant changes that can be attributed to genotype (G, accession), environment (E, temperature), and/or GxE interaction. Subsequently, we used the variance partitioning approach [38,39,41,42] to dissect and quantify the extent of the individual genotype and temperature effects on the phenotypic variation in more detail.

### Genotype, Environment, and GxE interaction analysis

Each phenotypic trait was subjected to a 2-factorial ANOVA to address which of the analyzed factors (G, E, GxE) had significant effects on the trait. Reaction norm plots for each phenotype are shown in Additional file 3. Each of the analyzed traits showed significant effects of genotype, environment (temperature) and GxE interaction (Additional file 15). Surprisingly, this included all juvenile and adult vegetative stages despite their seemingly uniform impression of temperature responses given by the Q_10_ values (Fig. 3a, b).

To assess genotype and temperature contributions in a more quantitative manner, we next used a variance partitioning approach [38,39,41,42]. Specifically, we calculated the index of phenotypic divergence (P_st_, [38]) at each analyzed temperature as a measure of genotype effects 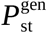 on the trait of interest (Additional file 16a). To complement this analysis, we also estimated the variation occurring across temperatures 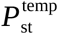 for each of the analyzed accessions (Additional file 16b), which enabled us to assess the temperature effect for the trait of interest for specific genotypes.

### Genotype effects

The 2-factorial ANOVA design of the GxE interaction analysis has shown that the genotype significantly affects variation of phenotypic traits. The variance partitioning index for genotype effects (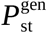) can extend this analysis by providing a quantitative assessment of the genotype contribution to variation at individual temperatures.

Individual 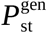 values showed highly variable patterns among the different traits and phenotype classes (Additional file 16a). Regardless of the individual temperature, mean genotype effects on developmental timing throughout the vegetative phase were generally very low (Fig. 4a), corroborating the impression gained from the analysis of Q_10_ values (Fig. 3). However, genotype effects on later stages of adult vegetative development seem to increase with higher temperatures (Additional file 16a), which may be the significant effect observed in the ANOVA-based GxE interaction assessment.

**Figure 4:**
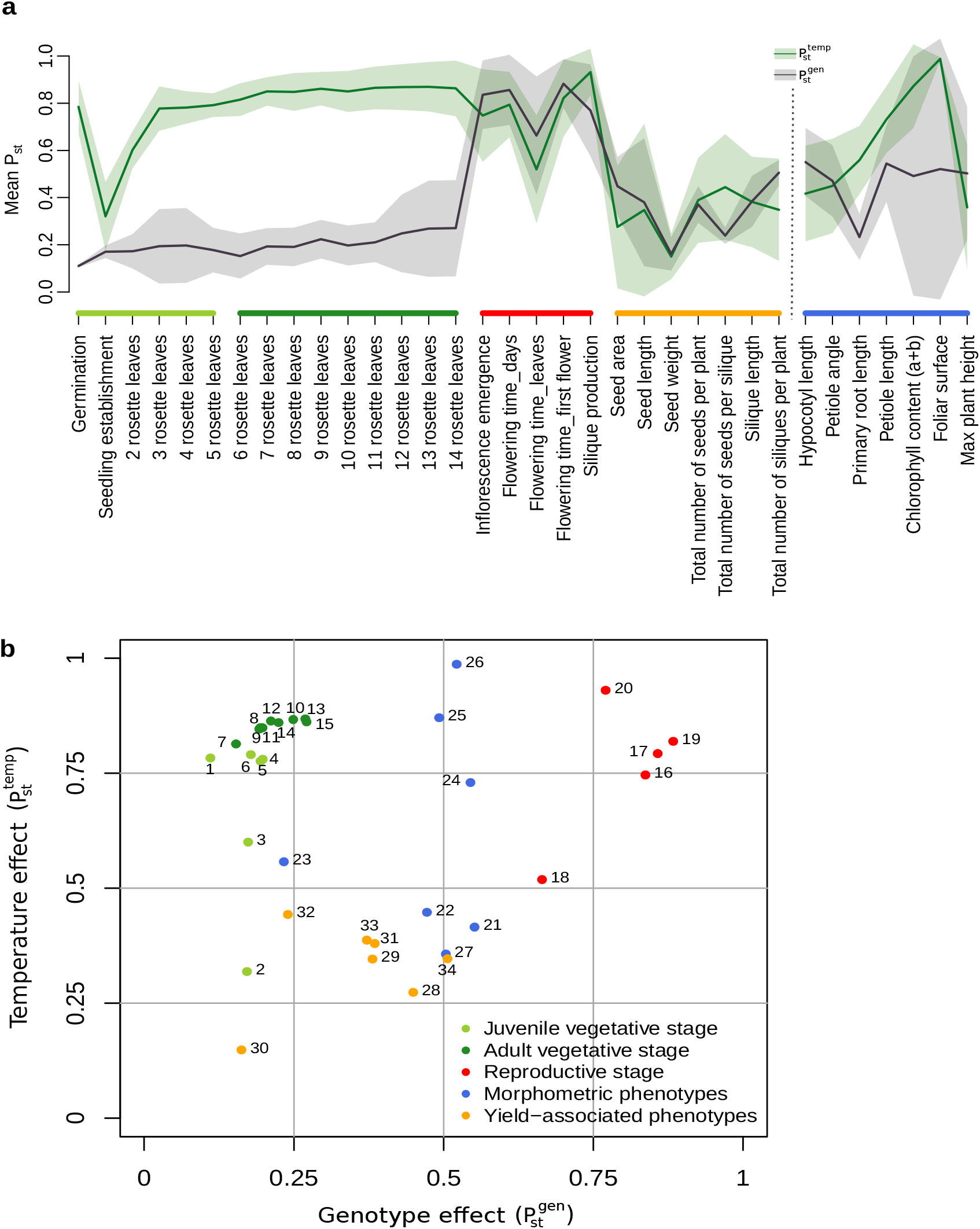
Genotype and temperature effects on phenotypic variation. (a) Genotype (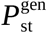, black) and temperature (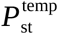, green) contribution to variation. Solid lines show mean P_st_ values and shaded areas indicate standard deviations. (b) Scatter plot of mean 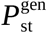 and 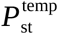 values over all temperatures and accessions, respectively. Phenotypes are color-coded according to the phenotype classes shown in Fig. 1 and described in Supporting Information Table S1. A heatmap of individual 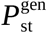 and 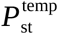 values and a scatter plot including standard deviations are shown in Additional file 16.

Similarly, strong genotype effects at higher temperatures were also observed for reproductive traits. Here, 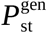 values at 16°C were already considerably higher than for vegetative growth stages and increased further with elevated temperatures (Additional file 16a). A contrasting pattern of decreasing genotype effects with an increase in temperatures was observed for total plant height indicating that here, natural variation in growth is higher at lower temperatures. Yield-associated phenotypes in general showed only low genotype effects on variation, indicating that under our experimental conditions variation in trait expression in this category is primarily affected by temperature (Fig. 4a).

Other phenotypes display rather differential or less gradual genotype effects among different temperatures. For example, the genotype impact on variation in hypocotyl and petiole length sharply increases from 24 to 28°C, indicating a certain buffering capacity or a threshold for natural variation.

In some cases, such as flowering time, a strong genotype effect seems to correlate also with a strong general temperature sensitivity as indicated by the high between-accessions variability in Q_10_ values (Fig. 4a and Fig. 3b). However, this does not seem to be a general principle. In case of root length, for example, low genotype i effects were observed (Fig. 4a, b), even though the phenotype in principle was highly sensitive to a change in ambient temperature (Fig. 3b).

### Temperature effects

We also used the variance partitioning approach to analyze the extent of the significant impact of temperature on phenotypic variation that was detected in the GxE interaction analysis (Additional file 15). Therefore, we calculated the index for temperature effects (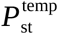) on the variation of phenotypic plasticity across all four temperatures within each of the ten accessions (Additional file 16b). While the 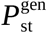 provided information on the genotype effect and thus, the overall natural variation of trait expression at different temperatures, the 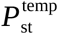 provides information primarily on the temperature-induced variability for each accession individually.

The heatmap representation of temperature effects (Additional file 16b) partially complements the genotype effect results. For example, variation in the timing of vegetative development was highly affected by temperature (high 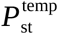), whereas 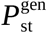 values were generally low (Fig. 4a, Additional file 16a, b). Interestingly, temperature effects in juvenile vegetative stages seemed to be lower (for seedling establishment and 2 rosette leave stage) than in later vegetative stages with the exception of germination which showed strong temperature effects in most accessions.

Many traits exhibit highly differential temperature effects among accessions in the sense of one accession demonstrating a particularly strong temperature effect on a specific trait, while another accession may show low to no temperature effects (e.g. chlorophyll content in Ler-1 vs. Bay-0). This is particularly obvious for yield-related traits such as total number of seeds per plant and silique as well as silique length. Here, temperature effects on phenotype variation were low for Col-0, C24 and Bay-0, whereas considerably higher 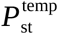 values were determined for the other accessions (Additional file 16b). Accessions which exhibit strong temperature effects on phenotypic variation may be interesting candidates for forward genetic approaches to identify the contributing molecular regulatory components.

### Comparison of temperature and genotype effects

As each phenotypic trait has been assigned a value for genotype and temperature effects, they can easily be compared to assess which of the two has a stronger influence on the phenotypic plasticity. To allow a direct comparison of effects, we compared mean values for 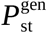 across all temperatures and 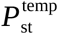 across all accessions (Fig. 4a, b).

Temperature effects on vegetative development showed a high, largely robust impact with little variance in 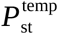 values, whereas genotype effects were generally low with diverging variances. Genotype effects peak at the transition to the reproductive phase and in some morphometric phenotypes. In general, morphometric parameters show high temperature and varying genotype effects. Phenotypes associated with late developmental stages were generally less affected by both factors indicating an overall buffering effect. Yet, variances in temperature effects tended to be high here, which may indicate genotype-specific thresholds for temperature effects (Fig. 4a, Additional file 16c). A scatter plot representation of mean 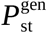 and 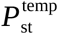 values for each trait allows further comparison of phenotypes according to the impact of both factors (Fig. 4b). While vegetative and reproductive phenotypes form tight clusters, morphometric phenotypes displayed a heterogenous pattern. In these traits, temperature responses seem to be affected by natural variation and may thus serve as candidate phenotypes for classic or quantitative forward genetic analyses.

Several yield-associated phenotypes such as total number of seeds, seed size and seed weight showed varying degrees of temperature sensitivity, likely caused by the partially distinct temperature effects on individual accessions (Fig.2b, Additional file 17).

The fundamental impact on temperature on the phenotypic responses is also reflected in the results of the principle component analysis (PCA). The PCA was performed on mean-centered and scaled data in order to allow integration of data with different scaling. PC1 which covered 50% of the observed variation, allowed a clear separation of samples via temperature (Fig. 5a). Here, the differentiation between 16 and 20 °C seems to be higher than the temperature changes from 20 to 24 °C and 24 to 28°C. PC2 explained ~16% of the variation and separated samples rather by genotype. Here, Rrs-7 and Got-7 showed a clear divergence from other genotypes. Again, this separation is already clear between 16 and 20°C whereas a further increase in temperature contributed little more to the separation.

**Figure 5:**
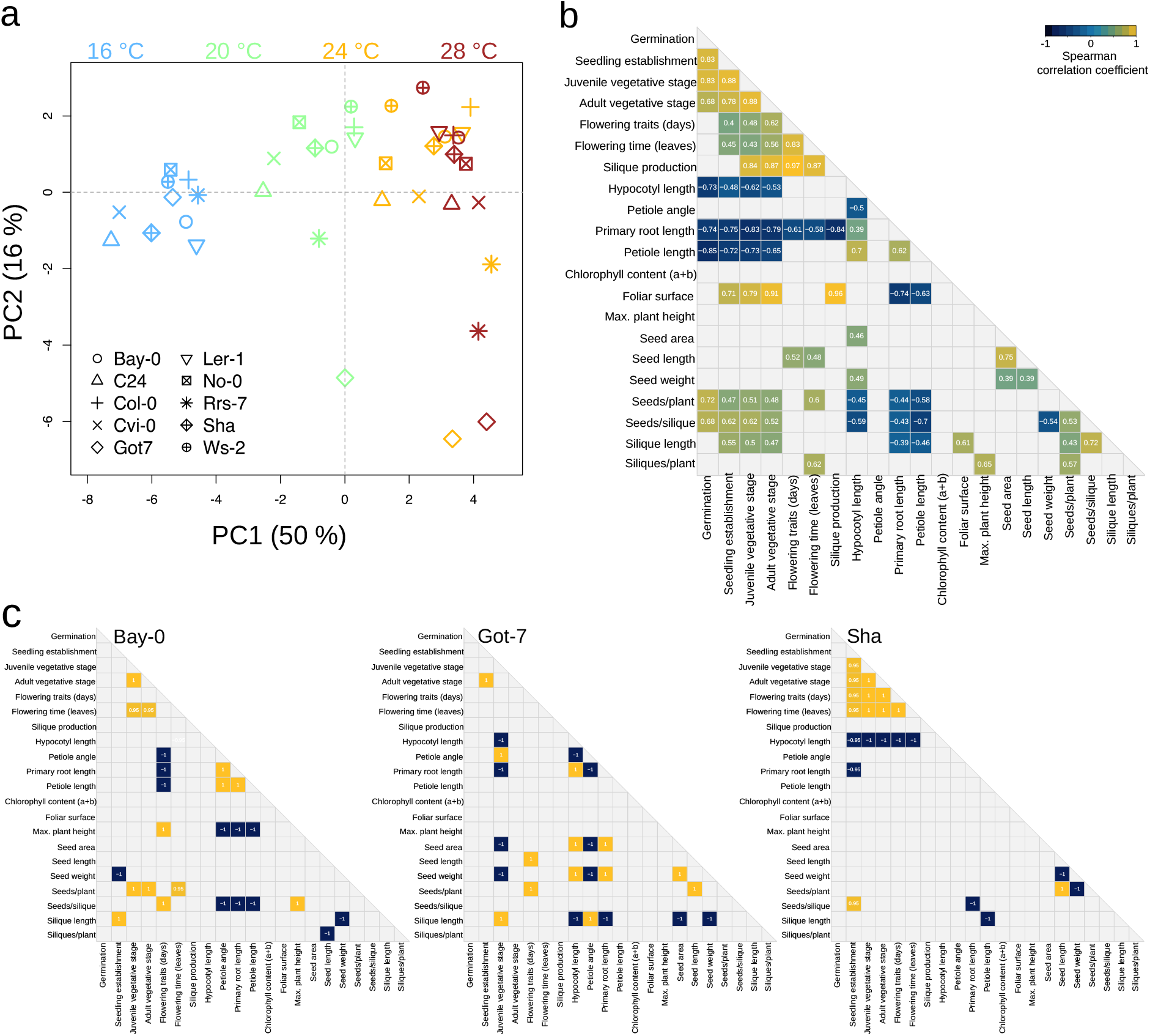
Principle component and correlation analyses. (a) Phenotypic data of all temperatures and genotypes were subjected to principle component analysis (PCA). (b-c) Correlation analysis of temperature responses among individual traits or trait groups of all analyzed genotypes (b) or in selected individual accessions (c). Spearmann correlation coefficients were tested for significance and coefficients with P < 0.05 and P < 0.1 are presented in (b) and (c), respectively. Phenotype correlations for all accessions individually are shown in Additional file 18.

### Correlation of phenotypic temperature responses

Finally, we analyzed putative correlations in temperature responses among different phenotypes to assess whether individual phenotype responses are indicative of temperature responses in general. As redundancies of individual phenotypes may bias the analyses several traits were combined in groups for further analyses (e.g. rosette development or flowering traits). We used the rank-based Spearman correlation coefficients for pairwise comparisons of averaged trait (group) values among all accessions to account for potential non-linear relationships and minimize outlier effects. As to be expected from the varying degrees of genotype and temperature effects on different traits, phenotypic correlations also varied considerably. To filter for robust correlations, only significant correlations (P < 0.05) were retained in the analysis (Fig. 5b).

High correlations were detected among traits within the vegetative stage of development (e.g. juvenile and adult vegetative stage), and among traits within the reproductive phase (e.g. flowering traits and the onset of silique production). In addition, temperature-induced reduction in foliar surface correlated strongly with the decrease in developmental time in vegetative and reproductive phases. Similarly, the reduction in developmental times and foliar surface were moderately correlated to the effect on several seed-associated traits (Fig. 5b).

Model temperature phenotypes such as petiole and hypocotyl length showed a positive correlation and were in turn correlated or inversely correlated with several other phenotypes or trait groups. However, temperature responses in primary root length under these experimental conditions showed an even more robust connection to many other traits. Mostly, these were inverse correlations with the exception of other seedling traits which were positively correlated with primary root lengths (Fig. 5b).

Due to the differential genotype effects on variation we also wondered whether individual genotypes may show different correlation patterns among phenotypic temperature responses. Calculation of Spearmann correlation coefficients for each individual accession is based on a maximum of four data points per phenotype or trait group which generally results in weaker interactions among samples. Thus, the P-value threshold was set to 0.1 in the analysis which retained only the strongest (inverse) correlations. Inspection of the correlation patterns reveals remarkable differences among accessions (Fig. 5c, Additional file 18). For instance, petiole lenght, angle and primary root length in Bay-0 were all inversely correlated with flowering time, plant height and the number of seeds/silique, whereas in Sha, only hypocotyl lengths showed an inverse correlation with developmental timing in vegetative and reproductive stages. Got-7 even showed unique correlation patterns among early growth responses with inverse correlations among petiole angles and hypocotyl and root lengths, respectively (Fig. 5c). In general, the diversity in correlation patterns may indicate differential capacities for temperature responses that result in differential activation or buffering and, thus, in different extents of physiological temperature impacts. Elucidation of the underlying mechanisms of differential temperature responses and adaptations may provide essential tools for the modulation of crop responses to elevated ambient temperatures.

### Discussion

Increased ambient temperatures have previously been shown to affect i thermomorphogenesis for selected “model” phenotypes. A systematic assessment of developmental and phenotypic plasticity across a complete life cycle has, to the best of our knowledge, been lacking so far. This study aims to provide such a solid base of temperature effects on plants by consecutive profiling of plant growth and development throughout a life cycle of *A. thaliana* grown in four different ambient temperatures. Furthermore, including several distinct *A. thaliana* accessions reduced I potential genotype-specific biases in the data and allowed the analysis of temperature and genotype effects on the variation observed in different phenotypic traits.

All of the 34 analyzed phenotypes were significantly affected by different growth temperatures, natural variation, and GxE interactions, illustrating the fundamental impact of ambient temperature on plant development and the high variability in responses among genotypes (Additional files 4–13, 15). The variance partitioning approach allowed the further dissection of phenotypes based on the extent of temperature and genotype effects. First, we identified phenotypes that were primarily affected by temperature and showed small genotype-induced variation. Second, we i identified phenotypes that additionally or even predominantly showed genotype effects on the observed phenotypic variation.

Developmental timing of juvenile and adult vegetative growth was significantly affected by genotype and temperature (Additional file 15). Yet, temperature was the dominant factor in the observed variation (Fig. 4a, 5a, Additional file 16). Genotype effects, albeit significant, were limited and mostly showed similar accelerations by increasing temperatures in all analyzed genotypes. This observation may be indicative for extensive thermodynamic effects on (conserved) regulatory mechanisms involved in this process. Indeed, thermomorphogenic responses are often speculated to be primarily caused by broad or general effects of free energy i changes on biochemical reactions (e.g. enzyme activities). The validity of the early proposed temperature coefficient (Q_10_) for plant development was demonstrated for germination rates and plant respiration [43,44]. The strong temperature effect on the acceleration of developmental timing throughout the vegetative phase, which was only weakly affected by genotypes supports this theory. When adopting the terms of “passive” and “active” temperature effects as proposed by [45], timing of vegetative development would represent a passive temperature response that might be caused by thermodynamic effects on metabolic rates and enzyme activities or on highly conserved signaling/response components.

On the other hand, phenotypes that show a high degree of genotype and temperature effects might rather be influenced by one or more specific genes that contribute to trait expression in a quantitative manner. As such, these phenotypes would represent “active” temperature effects [45]. However, the involvement of specific signaling elements does not necessarily exclude influences via thermodynamics. In fact, the recently described thermosensing via phyB acts via the promotion of phyB P_FR_ to P_R_ conversion in a temperature-promoted manner [18,19]. Natural variation in thermomorphogenic responses could be caused by polymorphisms in signaling or response genes ranging from alteration in gene sequence to expression level polymorphism [46]. As they may provide keys to altered temperature responses that could be utilized in specific breeding approaches, identification of such genes would be of high interest.

In fact, natural allelic variation in the circadian clock components *ELF3* and in the regulation of *GIGANTEA* have recently been shown to directly affect PIF4-mediated hypocotyl elongation in response to elevated temperatures [12,13,47]. Therefore, PIF4 and PIF4-regulating components could be important targets of adaptation to growth in higher ambient temperatures. PIF4 and ELF3 have been shown to be involved in both, temperature-induced hypocotyl elongation and the induction of flowering [12,13,20,48]. However, a lack of general correlation among seedling growth and flowering time responses may indicate that these processes are not universally regulated via the same components. Alternatively, the impact of these signaling components on diverse phenotypes may be more prominent for specific alleles which may be reflected by the diversity in correlation patterns among individual accessions (Fig. 5c, Additional file 18).

In general, the intraspecific diversity in phenotypic changes in response to elevated ambient temperatures argue against a general explanation of morphological and developmental changes due to passive thermodynamic effects.

Exploiting natural genetic variation to identify genes that are involved in the regulation of temperature effects on specific traits can provide new leads for plant breeding. The work presented here may inspire new approaches for temperature research in non-reference accessions as some temperature responses were much more pronounced in accessions other than Col-0 (Fig. 3b). Specific approaches will depend on the focus on either yield-or biomass-associated traits.

In conclusion, our work provides a map that allows the dissection of thermomorphogenesis in phenotypic traits that are either robustly affected by temperature or traits that are differentially affected by temperature among different accessions. While robust temperature-sensitive phenotypes might indeed be caused by thermodynamic acceleration of metabolism or highly conserved signaling events, natural genetic variation of temperature responses implicate the relevance of specific regulatory cascades that can be instrumental to future breeding approaches.

## Declarations

### Ethics approval and consent to participate

Not applicable

### Consent for publication

Not applicable

### Availability of data and material

The datasets analysed during the current study is available from the corresponding author on request.

### Authors' contributions

CI, MQ, and CD designed the research and experimental setup. CI, TP, JB and KD performed the phenotypic analyses and data collection. CI, YP and CD analyzed the data. YP, AG-D, and CD designed and performed statistical analyses. CI, YP, MQ, and CD interpreted data, prepared figures and wrote the manuscript.

## Acknowledgements

Not applicable

## Additional files

Additional file 1: Table of recorded phenotypes and association to phenotype classes

Additional file 2: Identity and geographic origin of analyzed *A. thaliana* accessions

Additional file 3: Reaction norm plots of each phenotype for each of the analyzed genotypes

Additional file 4: Summary of Col-0 thermomorphogenesis

Additional file 5: Summary of Bay-0 thermomorphogenesis

Additional file 6: Summary of C24 thermomorphogenesis

Additional file 7: Summary of Cvi-0 thermomorphogenesis

Additional file 8: Summary of Got-7 thermomorphogenesis

Additional file 9: Summary of Ler-1 thermomorphogenesis

Additional file 10: Summary of No-0 thermomorphogenesis

Additional file 11: Summary of Rrs-7 thermomorphogenesis

Additional file 12: Summary of Sha thermomorphogenesis

Additional file 13: Summary of Ws-2 thermomorphogenesis

Additional file 14: Natural variation in developmental timing (leaves vs. days)

Additional file 15: GxE interaction analysis results

Additional file 16: Detailed information on genotype and temperature effects on phenotypic variation

Additional file 17: Temperature effect on yield

Additional file 18: Correlations among temperature responses in individual accessions

## References

1. Quint M, Delker C, Franklin KA, Wigge PA, Halliday KJ, van Zanten M., Molecular and genetic control of plant thermomorphogenesis. Nat. Plants. 2016;2:15190.

2. Peng S, Huang J, Sheehy JE, Laza RC, Visperas RM, Zhong X, et al. Rice yields decline with higher night temperature from global warming. Proc. Natl. Acad. Sci. 2004;101:9971–5.

3. Moore FC, Lobell DB. The fingerprint of climate trends on European crop yields. Proc. Natl. Acad. Sci. U. S. A. 2015;112:2670–5.

4. IPCC. Climate change 2013: The physical science basis. Fifth assessment report. [Internet]. UNEP/WMO; Available from: http://www.ipcc.ch/report/ar5/wg1/.

5. Lobell DB, Gourdji SM. The Influence of Climate Change on Global Crop Productivity. Plant Physiol. 2012;160:1686–97.

6. Fitter AH, Fitter RSR. Rapid Changes in Flowering Time in British Plants. Science. 2002;296: 1689–91.

7. CaraDonna PJ, Iler AM, Inouye DW. Shifts in flowering phenology reshape a subalpine plant community. Proc. Natl. Acad. Sci. 2014;111: 4916–21.

8. Thuiller W, Lavorel S, Araújo MB, Sykes MT, Prentice IC. Climate change threats to plant diversity in Europe. Proc. Natl. Acad. Sci. 2005;102: 8245–50.

9. Gray WM, Östin A, Sandberg G, Romano CP, Estelle M. High temperature promotes auxin-mediated hypocotyl elongation in Arabidopsis. Proc. Natl. Acad. Sci. 1998;95: 7197–202.

10. Zanten M van, Voesenek LACJ, Peeters AJM, Millenaar FF. Hormone-and Light-Mediated Regulation of Heat-Induced Differential Petiole Growth in Arabidopsis. Plant Physiol. 2009;151: 1446–58..

11. Balasubramanian S, Sureshkumar S, Lempe J, Weigel D. Potent Induction of Arabidopsis thaliana Flowering by Elevated Growth Temperature. PLoS Genet. 2006;2: e106..

12. Raschke A, Ibañez C, Ullrich KK, Anwer MU, Becker S, Glöckner A, et al. Natural Variants of ELF3 Affect Thermomorphogenesis by Transcriptionally Modulating PIF4-Dependent Auxin Responses. BMC Plant Biol. 2015;15: 197..

13. Box MS, Huang BE, Domijan M, Jaeger KE, Khattak AK, Yoo SJ, et al. ELF3 Controls Thermoresponsive Growth in Arabidopsis. Curr. Biol. 2015;25: 194–9.

14. Zhu W, Ausin I, Seleznev A, Mendez-Vigo B, Pico FX, Sureshkumar S, et al. Natural Variation Identifies ICARUS1, a Universal Gene Required for Cell Proliferation and Growth at High Temperatures in Arabidopsis thaliana. PLoS Genet. 2015;11: e1005085.

15. Lutz U, Posé D, Pfeifer M, Gundlach H, Hagmann J, Wang C, et al. Modulation of Ambient Temperature-Dependent Flowering in Arabidopsis thaliana by Natural Variation of FLOWERING LOCUS M. PLoS Genet. 2015;11: e1005588.

16. Sanchez-Bermejo E, Balasubramanian S. Natural variation involving deletion alleles of FRIGIDA modulate temperature-sensitive flowering responses in Arabidopsis thaliana. Plant Cell Environ. 2016;39: 1353–65.

17. Ma D, Li X, Guo Y, Chu J, Fang S, Yan C, et al. Cryptochrome 1 interacts with PIF4 to regulate high temperature-mediated hypocotyl elongation in response to blue light. Proc. Natl. Acad. Sci. 2016;113: 224–9.

18. Jung J-H, Domijan M, Klose C, Biswas S, Ezer D, Gao M, et al. Phytochromes function as thermosensors in Arabidopsis. Science. 2016;354: 886–89.

19. Legris M, Klose C, Burgie ES, Costigliolo C, Neme M, Hiltbrunner A, et al. Phytochrome B integrates light and temperature signals in Arabidopsis. Science. 2016;354: 897–900.

20. Koini MA, Alvey L, Allen T, Tilley CA, Harberd NP, Whitelam GC, et al. High Temperature-Mediated Adaptations in Plant Architecture Require the bHLH Transcription Factor PIF4. Curr. Biol. 2009;19: 408–13.

21. Franklin KA, Lee SH, Patel D, Kumar SV, Spartz AK, Gu C, et al. Phytochrome-interacting factor 4 (PIF4) regulates auxin biosynthesis at high temperature. Proc. Natl. Acad. Sci. U. S. A. 2011;108: 20231–5.

22. Proveniers MCG, van Zanten M. High temperature acclimation through PIF4 signaling. Trends Plant Sci. 2013;18: 59–64.

23. Toledo-Ortiz G, Johansson H, Lee KP, Bou-Torrent J, Stewart K, Steel G, et al. The HY5-PIF Regulatory Module Coordinates Light and Temperature Control of Photosynthetic Gene Transcription. PLoS Genet. 2014;10: e1004416.

24. Delker C, Sonntag L, James GV, Janitza P, Ibañez C, Ziermann H, et al. The DET1-COP1-HY5 Pathway Constitutes a Multipurpose Signaling Module Regulating Plant Photomorphogenesis and Thermomorphogenesis. Cell Rep. 2014;9: 1983–9.

25. Gangappa SN, Kumar SV. DET1 and HY5 Control PIF4-Mediated Thermosensory Elongation Growth through Distinct Mechanisms. Cell Rep. 2017;18: 344–51.

26. McKhann HI, Camilleri C, Bérard A, Bataillon T, David JL, Reboud X, et al. Nested core collections maximizing genetic diversity in Arabidopsis thaliana. Plant J. 2004;38: 193–202.

27. Delker C, Pöschl Y, Raschke A, Ullrich K, Ettingshausen S, Hauptmann V, et al. Natural Variation of Transcriptional Auxin Response Networks in Arabidopsis thaliana. Plant Cell. 2010;22: 2184–200.

28. Scholl RL, May ST, Ware DH. Seed and Molecular Resources for Arabidopsis. Plant Physiol. 2000;124: 1477–80.

29. Lincoln C, Britton J, Estelle M. Growth and development of the axr1 mutants of Arabidopsis. Plant Cell. 1990;2: 1071–80.

30. ImageJ: http://imagej.nih.gov/ij/

31. RootDetection: http://www.labutils.de/rd.html

32. Boyes DC, Zayed AM, Ascenzi R, McCaskill AJ, Hoffman NE, Davis KR, et al. Growth stage-based phenotypic analysis of Arabidopsis: a model for high throughput functional genomics in plants. Plant Cell. 2001;13: 1499–510.

33. Porra RJ, Thompson WA, Kriedemann PE. Determination of accurate extinction coefficients and simultaneous equations for assaying chlorophylls a and b extracted with four different solvents: verification of the concentration of chlorophyll standards by atomic absorption spectroscopy. Biochim. Biophys. Acta BBA - Bioenerg. 1989;975: 384–94.

34. R Core Team. R: A Language and Environment for Statistical Computing [Internet]. Vienna, Austria: R Foundation for Statistical Computing; 2015: https://www.R-project.org

35. Nicotra AB, Atkin OK, Bonser SP, Davidson AM, Finnegan EJ, Mathesius U, et al. Plant phenotypic plasticity in a changing climate. Trends Plant Sci. 2010;15:684–92.

36. Whitman D, Agrawal A. What is Phenotypic Plasticity and Why is it Important? Phenotypic Plast. Insects. Science Publishers; 2009: http://dx.doi.org/10.1201/b10201-2

37. Loveys BR, Atkinson LJ, Sherlock DJ, Roberts RL, Fitter AH, Atkin OK. Thermal acclimation of leaf and root respiration: an investigation comparing inherently fast-and slow-growing plant species. Glob. Change Biol. 2003;9: 895–910.

38. Storz JF. Contrasting patterns of divergence in quantitative traits and neutral DNA markers: analysis of clinal variation. Mol. Ecol. 2002;11: 2537–2551.

39. Leinonen T, Cano JM, Mäkinen H, Merilä J. Contrasting patterns of body shape and neutral genetic divergence in marine and lake populations of threespine sticklebacks. J. Evol. Biol. 2006;19: 1803–1812.

40. NASC/ABRC. https://www.arabidopsis.org/abrc/catalog/natural_accession_9.html

41. Gay L, Neubauer G, Zagalska-Neubauer M, Pons J-M, Bell DA, Crochet P-A. Speciation with gene flow in the large white-headed gulls: does selection counterbalance introgression? Heredity. 2008;102: 133–146.

42. Whitlock MC. Evolutionary inference from QST. Mol. Ecol. 2008;17: 1885–1896.

43. Hegarty TW. Temperature Coefficient (Q_10_), Seed Germination and Other Biological Processes. Nature. 1973;243: 305–6.

44. Atkin OK, Tjoelker MG. Thermal acclimation and the dynamic response of plant respiration to temperature. Trends Plant Sci. 2003;8: 343–51.

45. Penfield S, MacGregor D. Temperature sensing in plants. In: Franklin K a, Wigge P a, editors. Temperature and Plant Development. John Wiley & Sons, Inc; 2014. p. 1–18.

46. Delker C, Quint M. Expression level polymorphisms: heritable traits shaping natural variation. Trends Plant Sci. 2011;16: 481–8.

47. de Montaigu A, Giakountis A, Rubin M, Tóth R, Cremer F, Sokolova V, et al. Natural diversity in daily rhythms of gene expression contributes to phenotypic variation. Proc. Natl. Acad. Sci. 2015;112: 905–10.

48. Kumar SV, Lucyshyn D, Jaeger KE, Alós E, Alvey E, Harberd NP, et al. Transcription factor PIF4 controls the thermosensory activation of flowering. Nature. 2012;484: 242–5.

